# Genetic background determines allele-specific *Pfkelch13*-mediated artemisinin tolerance and persistence in *Plasmodium falciparum*

**DOI:** 10.64898/2026.09.03.749253

**Authors:** Imran Ullah, Morgan C. Martin, Daouda Ndiaye, Sarah K. Volkman, Dyann F. Wirth

## Abstract

Artemisinin-based combination therapies are the frontline treatment for *Plasmodium falciparum* malaria, but their efficacy is threatened by Artemisinin partial resistance (ART^R^). ART^R^ in *Plasmodium falciparum* is mediated by *Pfkelch13* (*k13*) mutations but remains rare in West Africa. A critical question is whether short-term drug survival intrinsically predicts long-term evolutionary persistence. Here, using CRISPR-Cas9 editing, and nanopore-assisted allele tracking in a contemporary Senegalese isolate, we evaluated ART^R^ receptivity using the ring-stage survival assay (RSA), a proxy for clinical resistance. We demonstrate that ART^R^ survival and drug-free persistence are biologically separable traits. While the WHO-validated Asian I543T mutation conferred the RSA phenotype, it was rapidly counter-selected during drug-free growth. In contrast, emerging African alleles (R561H, M579I) and a regional novel variant (C473S) establish stable resistance without a competitive penalty. These findings reveal that a parasite’s genetic background acts as a strict evolutionary filter, uncoupling short-term tolerance from long-term persistence and establishing that genomic context determines allele-specific resistance trajectories.

## Introduction

Artemisinin-based combination therapies (ACTs) are the frontline treatments for *Plasmodium falciparum* malaria worldwide. Consequently, the evolution and spread of artemisinin (ART) partial resistance (ART^R^) constitutes a major threat to global malaria control. Clinically, ART^R^ is characterized by delayed parasite clearance following ACT treatment. At the molecular level, this phenotype is primarily driven by nonsynonymous mutations in the propeller domain of the *k13* gene^1,2^. To monitor this threat, the World Health Organization (WHO) relies on the molecular surveillance of validated *k13* markers as an early-warning system to guide global antimalarial policy^3,4^. However, this framework implicitly assumes that an allele conferring acute drug tolerance will remain competitively viable and spread through the population.

In contrast to resistance to other antimalarials where resistance is mediated by a single primary mutation, ART^R^ can be mediated by multiple individual mutations throughout the propeller domain of the protein. Mechanistically, *K13* functions at the parasite cytostome, and resistance mutations confer drug tolerance by reducing hemoglobin endocytosis during the early ring stage^5^. This limits the release of heme required to activate the artemisinin prodrug^6^. However, this survival strategy presents a metabolic challenge: hemoglobin digestion is the parasite’s primary source of amino acids for rapid intraerythrocytic development. Consequently, restricting endocytosis to survive artemisinin frequently incurs substantial developmental fitness costs. This physiological trade-off likely explains why identical *k13* alleles confer vastly different levels of resistance and incur different fitness penalties depending on the parasite broader genetic background^2,7,8^. For example, Southeast Asian parasite lineages successfully acquired secondary, epistatic mutations (in genes such as *ferredoxin, apicoplast ribosomal protein S10, multidrug resistance protein 2 and chloroquine resistance transporter*) that buffered the physiological costs of mutated *k13* prior to the emergence of widespread resistance^9^. The evolutionary landscape in Africa, however, appears distinctly different. Currently, Africa bears 95% of the global malaria morbidity and mortality; and while ART^R^ has emerged in Eastern and Southern Africa, via validated *k13* alleles^10,11^, these mutations remain strikingly rare in West Africa^12–14^. This geographic disparity raises a critical biological question: is the West African *P. falciparum* genome inherently refractory to *k13*-mediated ART^R^, or do certain alleles simply incur fitness costs too severe to allow their establishment in highly competitive, high-transmission settings?

To address this, we investigated whether a contemporary West African parasite genome is capable of expressing *k13*-mediated ART^R^ and critically, whether alleles differ in their ability to persist once introduced. Using CRISPR-Cas9 genome editing and a novel nanopore-assisted allele tracking method in a recent Senegalese clinical isolate, we tracked both the acute drug survival and long-term drug-free competitive fitness of four distinct *k13* variants. By directly comparing the WHO-validated Asian I543T mutation against emerging African alleles, we demonstrate that a parasite’s genetic background acts as a strict evolutionary filter that can genetically uncouple short-term drug tolerance from long-term epidemiological persistence.

## Results

### Nanopore-assisted phenotyping reveals I543T confers artemisinin survival

As a proof of principle, we introduced four *k13* propeller variants (I543T, R561H, M579I and C473S) into a well-characterized recent Senegalese clinical isolate (SenTh15.14)^15^, using CRISPR–Cas9 technology^16^ (**Supplementary Fig. 1-2**). Baseline susceptibility to standard partner drugs was confirmed via standard growth assays (**Supplementary Fig. 3**), while ART response was assessed by the RSA^17^. Clonal edited lines were successfully obtained for R561H, M579I, and C473S transfections. However, despite repeated attempts, we were unable to recover a clonal I543T-edited line. I543T has been associated with high RSA survival, reaching 58% in Cambodian parasite isolates and 29% in the Dd2 reference background^1,2^. To bypass this cloning bottleneck and enable phenotyping, we developed nanopore-assisted RSA to measure variant allele frequency (VAF) immediately before dihydroartemisinin (DHA) exposure (**Fig. 1a**). Defined WT to Mutant (WT:Mut) genomic DNA mixtures showed close agreement between expected and observed mutant fractions to validate the assay (**Supplementary Fig. 4**). Using this approach, we quantified I543T frequency in the surviving parasites at 72 hours. While I543T frequency declined in drug-free controls (DMSO), DHA exposure strongly enriched the I543T population compared to matched controls (mean VAF 75.1% ± 18.9% in DHA vs 52.9% ± 22.5% in DMSO; P = 0.0158, paired t-test) (**Fig 1b**). A parallel nanopore RSA performed on a bulk-edited R561H population showed completely stable allele frequencies across all conditions (**Supplementary Fig. 5)**. Furthermore, standard microscopy-based RSA of the I543T bulk population demonstrated measurable overall survival (2.23 ± 0.60%; **Fig. 1c**), exceeding the established 1% *in vitro* threshold for ART^R17^. Thus, I543T retains the ability to confer measurable RSA survival in this Senegalese background.

**Figure 1:**
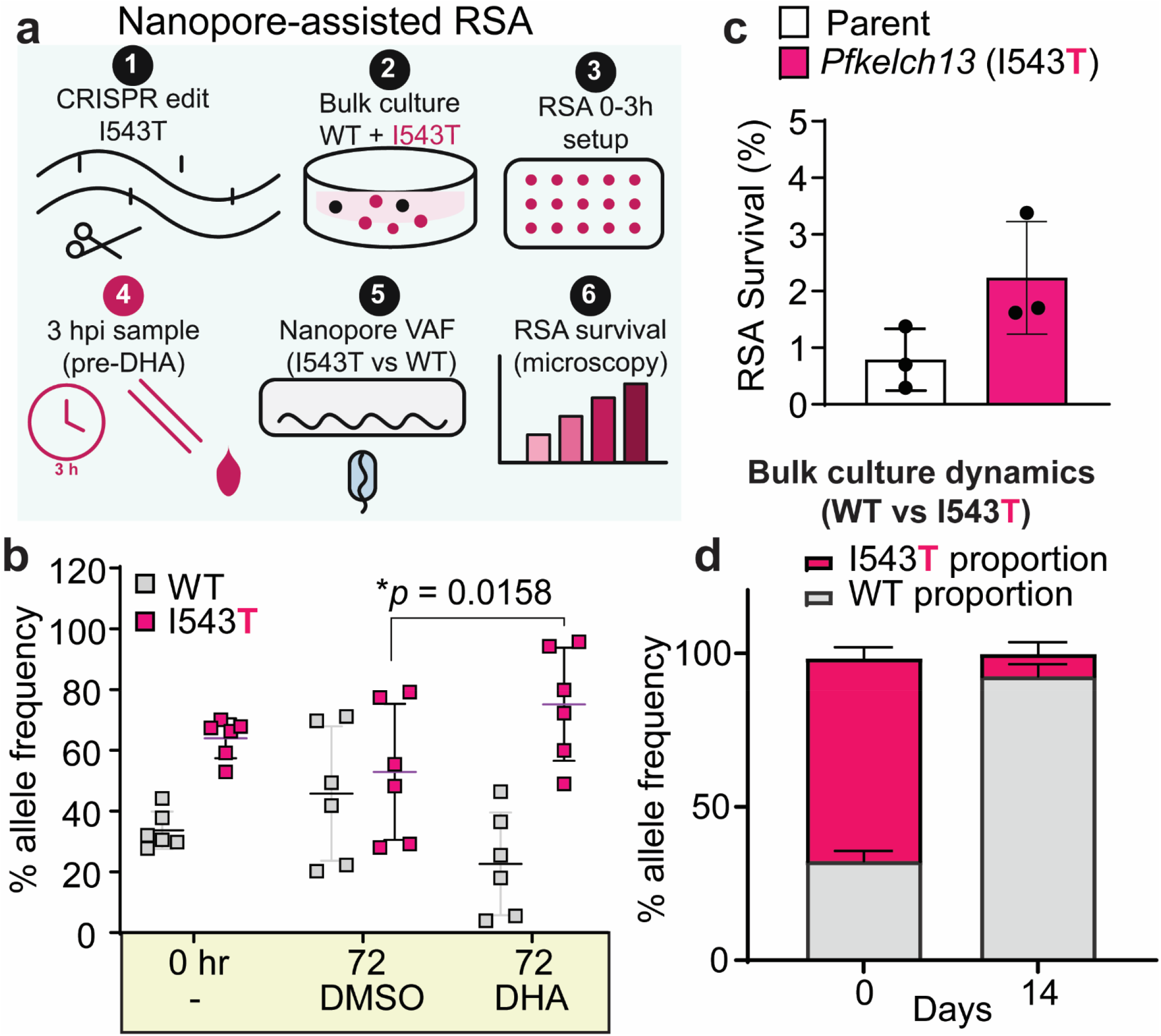
Nanopore-assisted RSA tracking and drug-free selection of bulk-edited I543T parasites in the SenTh15.14 background. **a**, Schematic of the nanopore-assisted RSA workflow for the bulk-edited I543T population. Bulk-edited I543T populations were expanded without clonal isolation. An aliquot was collected at 3 h post-invasion (hpi), before DHA exposure, for *k13* amplicon nanopore sequencing to quantify the variant allele frequency (VAF) in the starting RSA population, followed by sequencing of survivors at 72 h.x **b**, Nanopore-assisted tracking of I543T allele frequency. The VAF of the I543T mutation was quantified at 0 h and at 72 h following either a 6 h pulse with 700 nM DHA or treatment with DMSO vehicle. Data points represent technical replicates from three independent biological experiments; lines indicate the mean ± s.d. Statistical significance was determined using a two-tailed paired t-test comparing the averaged biological replicates from matched DHA- and DMSO-treated samples (P = 0.0158). **c**, Standard microscopy-based RSA survival (%) of the parental line and the bulk-edited I543T population (n = 3). Bars show the mean ± s.d. **d**, Drug-free bulk-culture dynamics. Two independent transfections were performed to generate separate bulk-edited I543T populations. Each population was maintained without drug selection and sampled at day 0 and 14. WT and mutant VAFs were quantified by *k13* amplicon nanopore sequencing. Stacked bars show the mean ± s.d. across the two biologically independent transfections. Source data are provided

### I543T imposes a severe competitive fitness cost during drug-free propagation

ART^R^ alleles can shape parasite evolution only if they survive the DHA pulse and persist when drug pressure is absent, a property not captured by RSA alone. We therefore tested whether introduced *k13* variants are stably maintained and competitively viable in the contemporary Senegalese SenTh15.14 background under drug-free conditions. Because clonal I543T parasites could not be recovered, we generated bulk-edited I543T populations in independent transfections and quantified *k13* allele frequencies during drug-free propagation. I543T was rapidly counter-selected: the mean I543T VAF declined precipitously from 66 ± 3.7% to just 7.2 ± 3.9% by day 14 without drug (**Fig. 1d**), whereas a bulk-edited R561H control remained stable over the same period (**Supplementary Fig. 6**). Together with the repeated failure to recover clonal I543T parasites, these longitudinal dynamics indicate a severe competitive fitness cost and strong negative selection against I543T in this Senegalese parasite background despite a measurable bulk RSA signal.

### Emerging African *k13* alleles establish stable resistance without a competitive penalty

We next sought to determine whether this severe fitness penalty was a universal feature of *k13* mutations in this isolate. Clonal edits of other alleles were readily recovered and showed strong RSA phenotypes in SenTh15.14 (**Fig. 1a-c**). The emerging African alleles R561H and M579I conferred high RSA survival (19.5 ± 5.9% and 27.6 ± 3.5%, respectively). The novel Senegal-associated allele C473S, which previously lacked functional characterization^18^, conferred an intermediate phenotype (3.96 ± 1.5%). Notably, these survival rates exceed those previously reported for the same alleles in other genetic backgrounds (R561H: 4–13% in Dd2/3D7/Cam3.II; M579I: 2–6% in CWX/3D7/UG659/UG815)^2,11,19,20^, consistent with strong genetic-background effects amplifying the magnitude of *k13*-mediated ring-stage survival^2,7,11^. To determine if these successful clones also lacked the competitive fitness penalty observed with I543T, we tracked them in head-to-head competitions initiated at nominal 1:1 mixtures with the parental line^21^. Over 30 days of drug-free culture, mutant VAF increased for R561H (mean paired increase +31.66 ± 2.29 percentage points), M579I (+31.45 ± 6.50), and C473S (+33.14 ± 13.07) (**Fig. 2d–f**). This demonstrates no overt replication or competitive deficit relative to the parental line under our *in vitro* conditions, allowing them to establish stable, persistent resistance phenotypes.

**Figure 2:**
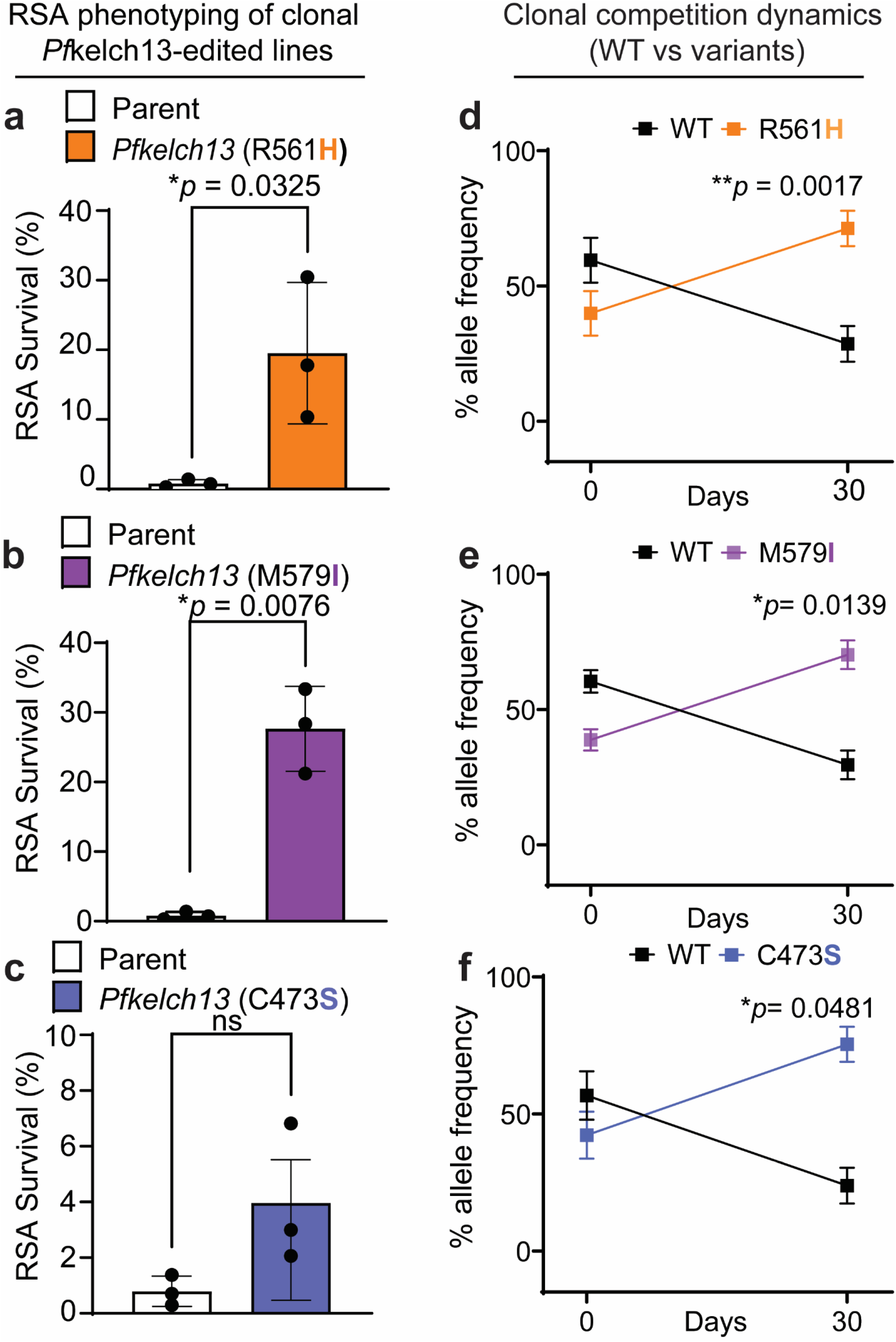
RSA phenotyping and clonal competition assays of *k13* variants in the SenTh15.14 background. **a–c**, RSA survival (%) for the SenTh15.14 parental line and clonal *k13*-edited lines carrying R561H (**a**), M579I (**b**), or C473S (**c**). Points indicate biologically independent experiments (n = 3); bars show the mean ± s.d. Statistical significance was assessed using a two-sided Kruskal–Wallis test with Dunn’s multiple-comparisons test versus the parental line; adjusted P values are shown. **d–f**, SenTh15.14 WT parasites were mixed at a nominal 1:1 ratio with isogenic clonal lines carrying R561H (**d**), M579I (**e**), or C473S (**f**). Mixed populations were maintained under standard drug-free *in vitro* culture conditions for 30 days. Cultures were sampled at days 0 and 30, and the variant allele frequency (VAF; percentage of nanopore reads supporting WT or the indicated *k13* allele) was quantified from *k13* amplicons. Black lines represent the WT proportion, and colored lines represent the corresponding mutant proportion. Points indicate the mean ± s.d. across biologically independent competition cultures (n = 3 per genotype). Statistical significance was determined using two-sided paired t-tests comparing mutant VAF at day 0 and day 30 within matched competition cultures: R561H, P = 0.0017; M579I, P = 0.0139; C473S, P = 0.0481. Source data are provided.

## Discussion

The emergence of ART^R^ is fundamentally an evolutionary filtering process. A resistance mutation must protect the parasite from acute drug toxicity, but it must also allow the parasite to replicate and transmit when the drug is absent. Our findings demonstrate that *k13*-mediated drug tolerance and drug-free persistence can be biologically uncoupled within a single contemporary West African parasite background. This highlights a critical limitation of relying solely on the RSA to predict the epidemiological threat of a given allele. An allele that promotes survival during a 6-hour *in vitro* drug pulse may still be an evolutionary dead-end if the parasite’s genetic background cannot tolerate the fitness consequences of that mutation.

Mechanistically, this uncoupling is likely tied to the parasite’s endocytic machinery and downstream stress-adaptation pathways. Previous studies have shown that *k13* variants mediate drug tolerance by downregulating hemoglobin endocytosis, which imposes a metabolic penalty via restricted amino acid availability^5,7^. Furthermore, surviving the resulting oxidative and proteostatic stress requires parasites to dramatically rewire their intraerythrocytic metabolism^22^. Our data are consistent with a model wherein this dual burden of nutrient starvation and altered stress-response imposes a physiological cost that the SenTh15.14 genome lacks the epistatic modifiers to buffer for the Asian I543T allele, leading to its rapid competitive exclusion in culture.

Importantly, our findings provide a critical nuance to previous studies of *P. falciparum* genetic backgrounds. While seminal work previously demonstrated that distinct geographic genomes (such as Southeast Asian versus African lineages) dictate the fitness costs of *k13* mutations^5^, our study reveals a highly selective paradigm within a single regional genome. We demonstrate that the contemporary West African genome is not universally refractory to ART resistance; rather, it acts as a stringent, allele-specific evolutionary sieve. It rapidly counter-selects the potent Asian I543T mutation, yet readily accommodates emerging African alleles (R561H, M579I) and a local novel variant (C473S) with no overt competitive penalty. This directly informs the current epidemiology of the region: highly deleterious alleles are unlikely to sweep through West Africa due to stringent fitness barriers, while these highly compatible African alleles are primed to spread.

Methodologically, our study provides a novel tool to investigate these evolutionary dynamics. In standard allelic replacement studies, deleterious edits are typically discarded as failed transfections because they cannot be cloned. Indeed, previous work in *Plasmodium berghei* demonstrated that several *k13* mutations could not be successfully introduced^23^. In *P. falciparum*, some *k13* alleles have reduced fitness when introduced into long-term culture-adapted parasites^24^. By developing a direct-from-culture, nanopore-assisted bulk RSA, we were able to quantitatively phenotype these unstable mutations. This approach allows the field to measure the exact boundary of parasite fitness and drug resistance, capturing phenotypes that are otherwise impossible to isolate clonally.

Moving forward, predicting the trajectory of ART resistance will require moving beyond single-locus paradigms. A priority next step is to identify secondary loci responsible for this background-specific buffering versus penalty. Where feasible, genetic crosses between an I543T-permissive background and SenTh15.14, coupled with selection under drug-free growth and/or defined DHA pulses, should enable mapping of modifiers that govern persistence. Ultimately, identifying the genomic contexts that permit resistance alleles to thrive will be essential for interpreting molecular surveillance data and predicting the true evolutionary trajectory of antimalarial resistance in Africa.

## Methods

### Parasite sampling, culture adaptation, and maintenance

The field isolate (ID: SenTh15.14) was extracted from a venous blood sample collected from a treatment-seeking patient in Thiès, Senegal in 2014. Ethical approval for sample collection was provided by the Senegal Ministry of Health and Social Action (Avis Protocol SEN1949) and the Harvard T.H. Chan School of Public Health Institutional Review Board (Protocol: 16330-CR). The isolate was culture-adapted as described by Brenneman et al. 2025 (denoted Th015.14)^15^. The line was whole genome sequenced pre- and post-culture adaptation also as described. The culture-adapted field isolate was cultured in RPMI 1640 medium supplemented with 10% Albumax and maintained by standard methods^15^.

### PCR amplification and sequencing

For *k13* amplification, a traditional or a modified (direct-from-culture) PCR setup was used. For the traditional PCR setup, genomic DNA (gDNA) served as PCR template. Blood pellets were collected from *in vitro* culture, and gDNA was extracted using the QIAmp DNA Blood Miniprep kit (Qiagen) following manufacturer’s instructions. For the direct-from-culture PCR setup, diluted culturing material served as PCR template. For sequencing confirmation and all experimentation outlined, a primer set was designed (IDT) to reliably produce ∼750bp amplicons covering a majority of the *k13* propeller domain coding sequence (CDS) (corresponding to residues 400-640) (*k13* Forward: CTTAGATAGGGATAGTGAGTTATTTAG; *k13* Reverse: GAACATAACATATTAGATTCCGTTGAAC) (primer sequences are also listed in Supplementary Table 1). All PCR reactions used Phusion High-Fidelity PCR Master Mix with HF Buffer (New England Biolabs, Inc.). All PCR amplification used the same thermocycling conditions as follows: 30s at 98°C; 35 cycles of three-step amplification at 98°C for 5s, 62°C for 15 s, and 72°C for 45s; and a final extension at 72°C for 5 min. PCR products were run on a 1% agarose gel for 15 min at 100V to confirm the expected size. PCR products were then purified using the Zymogen DNA cleanup kit, following the manufacturer’s instructions. Purified PCR products were submitted for Sanger sequencing (Genewiz) or for Oxford-Nanopore sequencing (Plasmidsaurus). For Sanger-sequenced PCR products, all chromatograms were analyzed against *k13* WT sequence covering the same region. For Nanopore-sequenced PCR products, all per-base data were analyzed to confirm *k13*-WT or mutant sequences; for mixed samples, the WT vs mutant (WT:MUT) proportions were calculated from the number of WT or MUT base calls divided by the total number of reads at target single-nucleotide polymorphism (SNP) positions. An average of approximately 1300 reads were obtained per PCR product sample.

### Gene editing of *k13* locus

#### Plasmid constructs

A two-plasmid strategy was used to express Cas9, the guide RNA (gRNA), and a donor repair template for homology-directed repair (HDR). Cas9 and the gRNA were co-expressed from an optimized derivative of the pDC2 CRISPR-Cas9 plasmid, pDC2-Cas9-U6-version2-hDHFR-noRep20, based on the system previously described^16^. The plasmid contains a *Plasmodium falciparum* U6 promoter for gRNA expression, a human dihydrofolate reductase (hDHFR) cassette conferring resistance to WR99210 for parasite selection, and an ampicillin-resistance marker for bacterial selection during plasmid propagation. Guide RNAs were designed using Benchling (Benchling, San Francisco, CA, USA). Guide oligonucleotides were annealed and ligated into the *BbsI*-digested Cas9 plasmid. The donor repair template was cloned into a pUC19 donor plasmid, which also contains an ampicillin-resistance marker for bacterial selection. Donor DNA was designed using the *k13* sequence from *P. falciparum* 3D7 (PF3D7_1343700) as the reference template. Each donor construct consisted of approximately 500 bp 5′ (upstream) and 3′ (downstream) homology arms flanking the target mutation. A central 10–20 bp engineered region encoded the desired nonsynonymous SNP together with synonymous shield mutations that disrupt gRNA/Cas9 recognition following HDR-mediated integration, thereby preventing re-cleavage of the edited locus. *SalI* and *XbaI* restriction sites were incorporated at the ends of the homology arms to facilitate cloning into the pUC19 donor backbone. In this two-plasmid CRISPR-Cas9 strategy, Cas9 introduces a double-stranded break at the target locus, and the donor template directs homology-directed repair, resulting in precise integration of the desired mutation into the parental genome (**Supplementary Fig. 1a**). The *Pf*Kelch13 domain architecture and the locations of edited residues are shown in **Supplementary Fig. 1b**. Correct assembly of both Cas9-gRNA and donor plasmids was confirmed by Sanger sequencing prior to parasite transfection. All gRNA oligonucleotides and mutagenesis primers (Integrated DNA Technologies, Coralville, IA, USA) used in this study are listed in **Supplementary Table 1**.

### Parasite culture, transfection, and cloning

*P. falciparum* SenTh15.14 WT asexual blood-stage parasites were cultured in O+ human red blood cells (RBCs) as previously described^15^. Upon reaching >3% parasitemia, consisting predominantly of ring-stage parasites, a sorbitol-synchronized 10 mL culture was prepared for transfection, using 2 mL of culture per plasmid DNA preparation. Prior to transfection, 50 μg of each plasmid (Cas9-gRNA and pUC19 donor) was precipitated in Cytomix. Plasmid DNA was electroporated into 80 μL packed RBCs using a Bio-Rad Gene Pulser (0.31 kV, 960 μF; time constant 8–11 ms) as previously described^8,21,25^. Drug selection was initiated 10–16 h after transfection by adding 5 nM WR99210 in RPMI-Albumax medium. Drug-containing medium was replaced daily for 3 days, after which cultures were maintained in drug-free medium with daily medium changes for an additional 3 days and every other day thereafter until parasites were detected by light microscopy. Transfectant parasites were typically detected 3–4 weeks after transfection. Following parasite recovery, gDNA was extracted once cultures reached approximately 1% parasitemia. Alternatively, bulk transfectants were screened by blood-direct PCR followed by Sanger sequencing to confirm successful genome editing. Confirmation of the edited *k13* alleles by Sanger sequencing (clonal R561H, M579I and C473S) and nanopore amplicon sequencing (bulk I543T) is shown in **Supplementary Fig. 2**. Successfully edited bulk populations were subsequently subjected to limiting dilution cloning to obtain clonal parasite lines

### Limiting dilution cloning

Bulk transfectant populations generated from the four engineered SenTh15.14 *k13* editing experiments were subjected to limiting dilution cloning to isolate clonal parasite populations. Serial dilutions of bulk transfectant cultures were distributed into 96-well plates and maintained at 2% hematocrit (HCT) with routine medium changes and culture expansion as required^8,21,25^. Wells were monitored weekly for parasite growth by Giemsa-stained thick blood smear microscopy. Upon detection of parasitemia, material from individual positive wells was used as template for PCR amplification of the *Pfkelch13* locus, followed by Sanger sequencing to confirm the presence of the intended mutations. Clonal parasite lines carrying the R561H, M579I, and C473S edits were successfully recovered. In contrast, despite successful Sanger confirmation of the I543T-edited bulk transfectant population, all clonal candidates recovered from this population retained the WT *k13* allele.

### Ring-stage Survival Assay (RSA)

The standard RSA was performed on early ring-stage parasites^8,17,21,26^. Briefly, a 1–2% late-stage schizont culture was enriched using a Percoll gradient prior to plating at 2% hematocrit and incubation at 37°C for 3 h to allow invasion. Following invasion, newly formed rings were synchronized with sorbitol, adjusted to 1% parasitemia at 2% HCT, and plated in triplicate for each treatment condition (200 μL per well in 96-well plates). The 0–3 h post-invasion ring-stage parasites were exposed to either 700 nM DHA or DMSO as a vehicle control for 6 h, after which the drug-containing medium was removed. At 72 h after initiation of drug treatment, parasitemia was quantified by Giemsa-stained thin blood smear microscopy, with 10,000 red blood cells counted per condition. Ring-stage survival was calculated as the percentage of viable DHA-treated parasites relative to DMSO-treated controls. An RSA survival value >1% was considered indicative of artemisinin (ART) partial resistance. Three biological replicates were performed for each parasite line, including WT controls and one clonal line per edited parasite population. Two independent smear counts were performed per condition, and RSA survival values are reported as mean ± SD (standard deviation).

### Concentration–response assays

All EC_50_ values were estimated using the standard 72 h Malaria SYBR Green I fluorescence assay^8,21,25–30^. Synchronized ring-stage parasites (1% starting parasitemia and 1% hematocrit) were cultured for 72 h at 37°C in 384-well plates containing serial dilutions of dihydroartemisinin (DHA; 0.05–500 nM), chloroquine (1–10,000 nM), monodesethyl-amodiaquine (1–10,000 nM), lumefantrine (0.05–500 nM), and piperaquine (0.05–1,000 nM). All compounds were resuspended in dimethyl sulfoxide (DMSO), with the exception of chloroquine, which was prepared in 0.1% Triton X-100 in water, and dispensed using an HP D300 Digital Dispenser. Concentration-response curves were generated using 24-point dilution series for DHA and 12-point dilution series for the remaining compounds, centered around the expected EC_50_ values for each drug. Each condition was tested in technical triplicate, and each plate included a DHA kill control and a no-drug growth control. Parasite viability was assessed after 72 h by SYBR Green I (Lonza) staining of parasite DNA. Fluorescence was measured using a SpectraMax M5 plate reader (Molecular Devices) with excitation at 494 nm and emission at 530 nm. Raw fluorescence values were normalized relative to the no-drug growth control, and EC_50_ values were calculated using GraphPad Prism 10 (GraphPad Software) by nonlinear regression using a log(inhibitor) vs response model with a four-parameter variable-slope curve fit. Dose-response curves and mean EC_50_ values (nM ± SD, standard deviation) are reported. Three independent biological replicates were performed for each parasite line and drug. Dose–response curves are shown in **Supplementary Fig. 3** and EC_50_ values are summarized in **Supplementary Table 2**.

### Nanopore quantification of known mixed WT:MUT ratios

SenTh15.14_WT and clonal SenTh15.14_R561H parasites were confirmed by Sanger sequencing prior to use. Genomic DNA from the two parasite lines was combined at five predefined WT:MUT ratios: 0:100, 25:75, 50:50, 75:25, and 100:0. The desired ratios were generated by mixing genomic DNA from each line based on DNA concentrations measured by Qubit fluorometric quantification. Each mixture was used as template for PCR amplification, and resulting amplicons were subjected to Nanopore sequencing as previously described^21^. Each WT:MUT mixture was independently prepared three times, with two technical PCR replicates performed per ratio. Per-base coverage at the codon encoding R561 was extracted from Nanopore sequencing data to determine the proportion of WT and MUT reads at the target position. The WT:MUT proportions determined by Nanopore sequencing for each predefined ratio are reported as mean (%) ± SD (standard deviation).

### Nanopore-assisted Ring-stage Survival Assay (RSA)

The standard RSA 0-3h was performed on ring-stage parasites. Briefly, late-stage schizont bulk transfectant culture was enriched using a Percoll gradient prior to plating at 2% hematocrit and incubation at 37°C for 3 h to allow invasion. Following invasion, sorbitol-synchronized rings were adjusted to 1% parasitemia at 2% HCT and plated in triplicate for each treatment condition (200 μL per well in 96-well plates). To determine the genetic composition of bulk parasite populations used for RSA, we developed a Nanopore-assisted allele quantification approach coupled to the standard RSA workflow. At 3 h post-invasion, immediately before DHA exposure, a small aliquot of culture was collected for direct-from-culture PCR amplification of the *k13* locus. The resulting amplicons represented the RSA starting (0 h) bulk population and were used to quantify WT:MUT allele proportions by Nanopore sequencing. The remaining culture was subjected to the standard RSA procedure, with 0–3 hpi ring-stage parasites exposed to either 700 nM DHA or DMSO vehicle control for 6 h, followed by removal of drug-containing medium. At 72 h after initiation of drug treatment, parasite survival was assessed by Giemsa-stained thin blood smear microscopy, with 10,000 RBCs counted per condition. RSA survival was calculated as the percentage of viable DHA-treated parasites relative to DMSO-treated controls. An RSA survival value >1% was considered indicative of ART partial resistance. Three biological replicates were performed for each bulk line, with two independent smear counts per condition, and RSA survival values are reported as mean ± SD (standard deviation). The 3 hpi PCR products were subjected to Nanopore sequencing, and per-base coverage at the codons encoding R561 or I543 was extracted to determine WT: MUT proportions.

### Long-term competitive fitness assay with Nanopore sequencing

SenTh15.14 parental WT parasites were mixed with each of the three clonal *k13* mutant lines (R561H, M579I, and C473S) at an approximately 1:1 starting ratio and maintained under standard *in vitro* culture conditions for 30 days. Long-term competitive fitness assays were performed with routine culture maintenance, including medium changes, culture splits, and monitoring of parasite viability. Samples were collected at Day 0 and Day 30, and genomic DNA was extracted from parasite blood pellets. The *k13* locus was amplified and subjected to Oxford Nanopore sequencing to quantify the relative abundance of WT and mutant alleles, as we have previously described^21^. Relative allele frequencies were determined at each time point, and changes in mutant allele abundance over the course of the competition were used to assess the relative competitive fitness of each *k13* allele in the SenTh15.14 genetic background. Three biological replicates were performed for each competition, with two technical replicates analyzed per sample.

### Allele frequency dynamics of *k13* edited bulk populations

The SenTh15.14_I543T allele was not stably maintained during culture, preventing recovery of a stable clonal I543T parasite line. Therefore, freshly generated I543T bulk transfectant populations from independent transfections were used for allele-frequency tracking experiments. Sanger-confirmed SenTh15.14_I543T and SenTh15.14_R561H (used as a control) bulk transfectant populations were maintained under standard *in vitro* culture conditions in the absence of drug selection for two weeks. To monitor changes in *k13* allele composition over time, a small aliquot of culture was collected from each population at weekly intervals for direct-from-culture PCR amplification of the *k13* locus. Bulk populations from two independent transfections per line were analyzed, resulting in two biological replicates with two technical PCR replicates per weekly collection. PCR products were subjected to Nanopore sequencing, and per-base coverage data were analyzed to quantify WT: MUT allele proportions. Changes in WT:MUT proportions over time were reported as mean percentage ± SD.

### Statistical analysis

All statistical analyses were performed using GraphPad Prism 10 (GraphPad Software, San Diego, CA, USA). Biological replicates were defined as independent experiments performed using independently maintained parasite cultures, independently generated edited parasite populations, or independently initiated competition assays, as specified for each experiment. Technical replicates were averaged prior to statistical analysis and were not treated as independent observations. Unless otherwise stated, data are presented as mean ± SD. All statistical tests were two-sided, and P values <0.05 were considered statistically significant. For ring-stage survival assays (RSA), survival was calculated for each biological replicate as the percentage of viable DHA-treated parasites relative to the corresponding DMSO-treated control. RSA survival values across parasite lines were compared using a Kruskal–Wallis test followed by Dunn’s multiple-comparisons test with the SenTh15.14 WT parental line as the reference group. For two-group RSA comparisons (e.g., WT vs bulk-edited I543T), a two-sided Mann–Whitney U test was used. For concentration–response assays, growth inhibition curves were fit independently for each biological replicate using nonlinear regression with a four-parameter variable-slope model (log[inhibitor] vs response) to estimate EC_50_ values. For each compound, EC_50_ values across parasite lines were compared using one-way ANOVA (comparisons performed relative to the SenTh15.14 WT parental line). For long-term head-to-head competition assays, *k13* variant allele frequency (VAF) was quantified from Oxford Nanopore sequencing as the proportion of reads supporting the mutant allele at the edited nucleotide position. Mutant VAF at Day 0 and Day 30 from the same competition culture were treated as paired observations and compared using a two-sided paired t-test. For validation of Nanopore-based allele quantification using predefined WT:MUT genomic DNA mixtures, the relationship between expected and observed mutant allele fractions was evaluated using simple linear regression (least squares); regression parameters (slope, intercept, R^2^, and 95% CI where applicable) are reported. For allele-frequency tracking in bulk-edited populations (e.g., I543T stability in drug-free culture), WT and mutant allele frequencies were quantified longitudinally by Nanopore sequencing and summarized descriptively as mean (%) ± SD across biological replicates.

## Author contributions

Conceptualization: I.U., S.K.V., D.F.W.; Methodology: I.U., M.C.M.; Investigation: M.C.M.; Formal analysis: I.U., M.C.M.; Resources: D.N.; Writing—original draft: I.U.; Writing—review and editing: D.N.; S.K.V., M.C.M., D.F.W., I.U.; Funding acquisition: D.F.W., S.K.V.; Supervision: I.U., D.F.W.

## Funding

This work was supported by the Bill & Melinda Gates Foundation (grant numbers INV-003442 and INV-019032 to D.F.W., and INV-049909 and INV-101391 to S.K.V.) and the National Institutes of Health (grant number R01 AI099105 to D.F.W.).

## Competing interests

The authors declare no competing interests.

## Data availability

Source data for all figures are provided with this manuscript. Additional data generated during this study are available from the corresponding author upon reasonable request.

## Notes

### Competing Interest Statement

The authors have declared no competing interest.

