## Supplemental Information for "Genetic background determines allele-specific *Pfkelch13*-mediated artemisinin tolerance and persistence in *Plasmodium falciparum*"

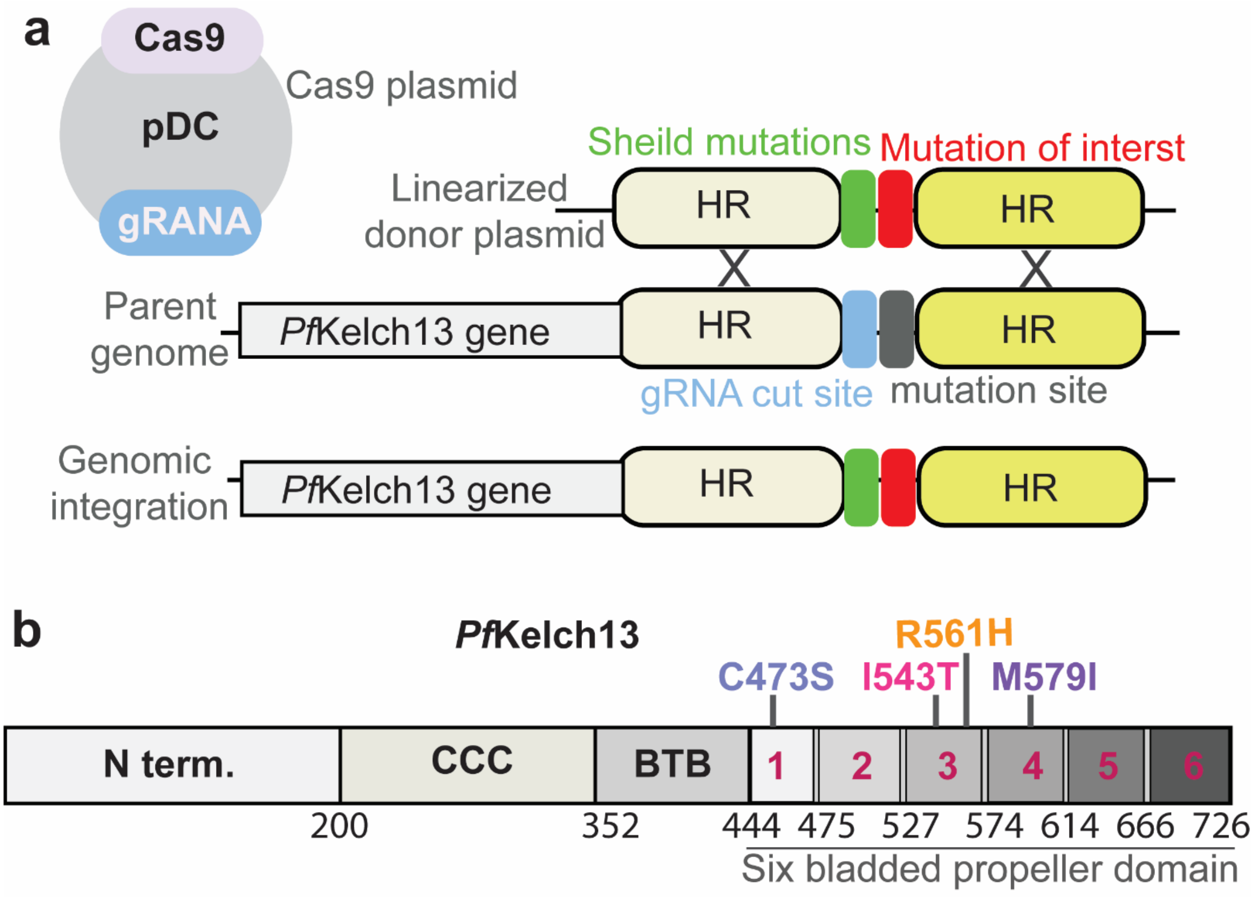


**Supplementary Fig. 1: CRISPR–Cas9 editing strategy and domain organization of *Pf*Kelch13.**

(**a**) Schematic of the plasmid used to introduce point mutations into the *Plasmodium falciparum Pfkelch13* locus. The Cas9 expression plasmid (Cas9, pDC) and the guide RNA (gRNA) expression cassette target a gRNA cut site within the endogenous *Pfkelch13* gene in the parent genome. A linearized donor plasmid carries two homology regions (HR) flanking the edited sequence. The donor insert contains (i) the desired mutation of interest (red) and (ii) additional synonymous "shielding" mutations (green) that disrupt the protospacer and/or PAM sequence without altering the encoded amino acids, thereby preventing re-cleavage by Cas9 after repair. Following double-strand break formation at the gRNA cut site and homology-directed repair using the donor plasmid, the edited allele is integrated at the native *Pfkelch13* locus, yielding a parasite line in which the mutation of interest and its associated shielding mutations are fixed in the genome.

(**b**) Linear map of the *Pf*Kelch13 showing the N-terminal region (N term.), the coiled-coil–containing (CCC) region, and the BTB/POZ domain, followed by the six-bladed β-propeller domain. Amino-acid positions corresponding to the boundaries between regions are indicated below the diagram. The six blades of the propeller domain are numbered (1–6), with the approximate residue ranges shown beneath each blade. The specific residues mutated in this study (C473**S**, I543**T**, R561**H**, and M579**I**) are indicated above their corresponding locations in the propeller domain.

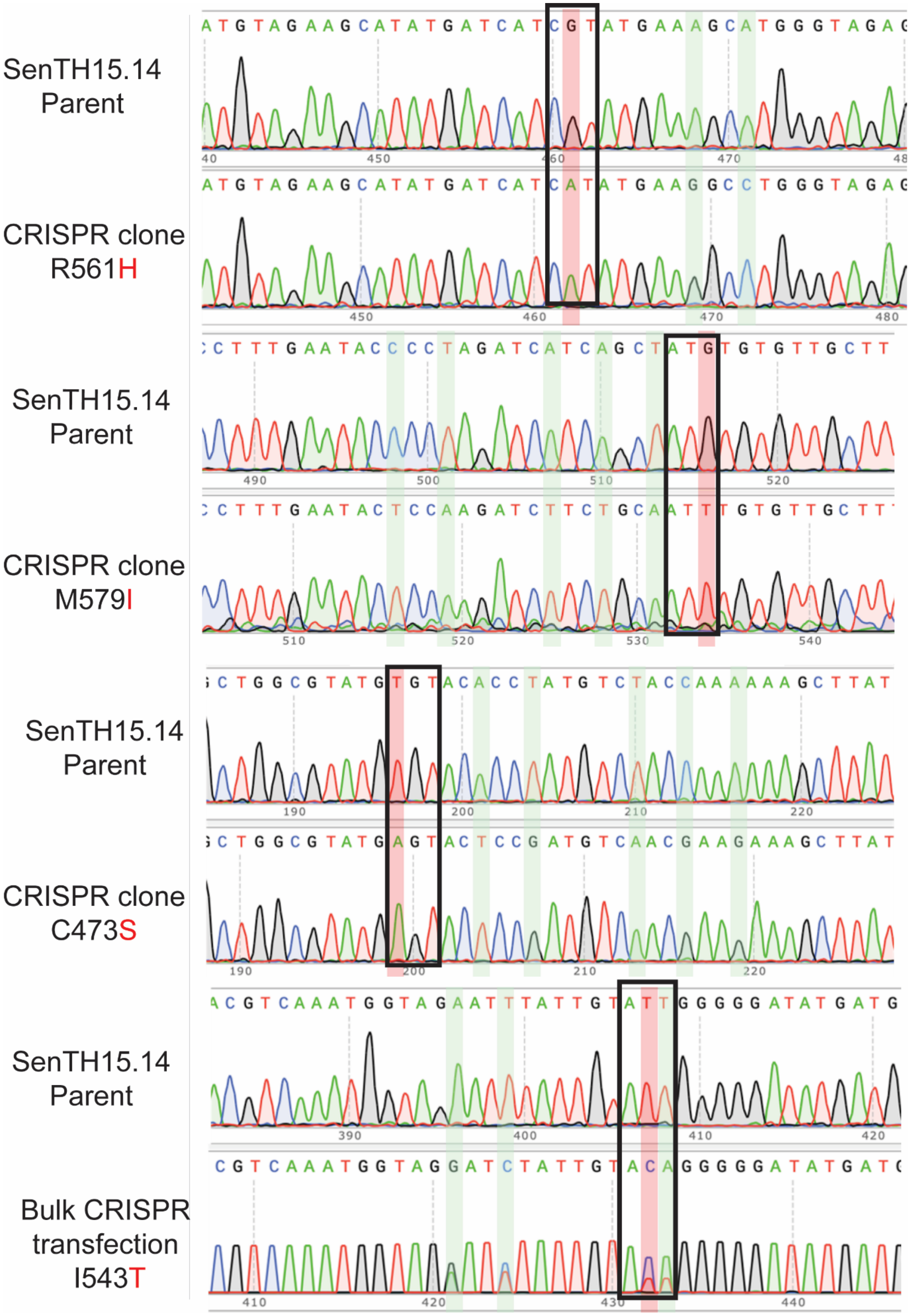
SenTH15.14 Parent
CRISPR clone R561H
SenTH15.14 Parent
CRISPR clone M579I
SenTH15.14 Parent
CRISPR clone C473S
SenTH15.14 Parent
Bulk CRISPR transfection I543T

**Supplementary Fig. 2. Sequencing confirmation of CRISPR–Cas9 editing at the endogenous *Pfkelch13* locus in the SenTH15.14 background.** Representative sequencing traces are shown for the SenTH15.14 WT parent and the corresponding clonal CRISPR-edited lines carrying the *Pfkelch13* mutations R561H, M579I, and C473S. The targeted codons are indicated by red-shaded vertical bands, and linked synonymous shielding substitutions introduced to disrupt the gRNA protospacer and/or PAM without altering the encoded amino acid are indicated by green-shaded bands. The engineered mutant codons are CAT for R561H, ATT for M579I, and AGT for C473S. For the I543T edit, sequence evidence from bulk-edited parasites was obtained by Nanopore amplicon sequencing, as indicated; the engineered mutant codon is ACA. The amino acid substitutions are boxed. Compared with the WT parent, the clonal edited lines show clean base calls at the targeted positions together with the linked shielding substitutions, consistent with precise allelic replacement at the endogenous *Pfkelch13* locus. The bulk I543T sample reflects a heterogeneous population containing both WT and edited alleles.

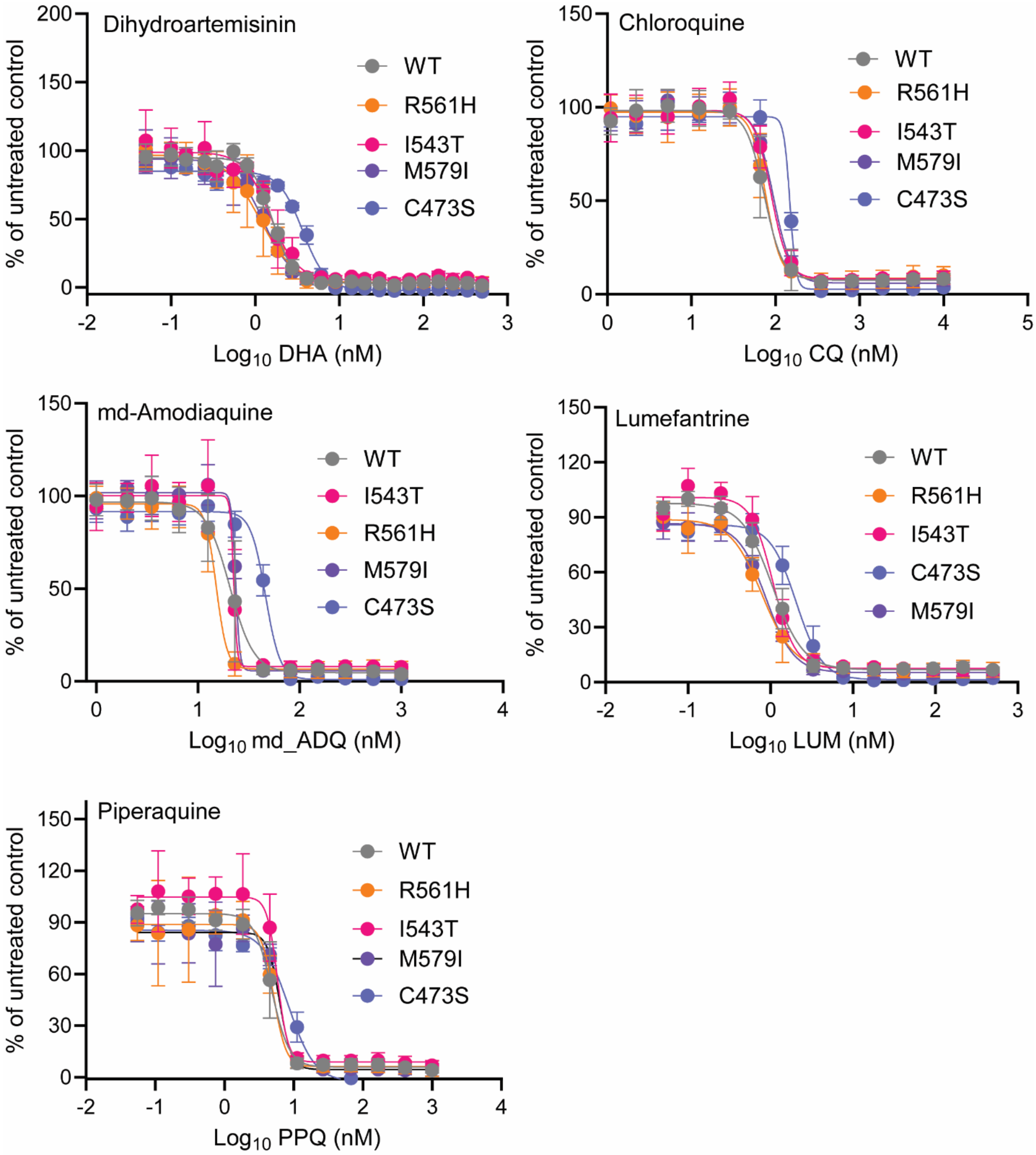


**Supplementary Fig. 3:** ***In vitro*** **dose–response ($EC_{50}$) curves for antimalarial drugs in SenTH15.14 WT and isogenic** ***Pfkelch13*****-edited lines**. Growth inhibition dose–response curves are shown for dihydroartemisinin (DHA), chloroquine (CQ), monodesethylamodiaquine (mdADQ), lumefantrine (LUM), and piperaquine (PPQ) against the SenTH15.14 WT parent and CRISPR-engineered *Pfkelch13* mutant lines (R561H, I543T, M579I, and C473S), all in the same genetic background. Drug concentrations are plotted as log10-transformed nanomolar (nM) values on the x-axis, and responses are normalized to the untreated control (% of untreated control) on the yaxis. Symbols show the mean response and error bars indicate ± SD from three independent biological replicates; solid lines represent the best-fit nonlinear regression for each parasite line (fit model as described in Methods). Corresponding $EC_{50}$ values derived from these fits are summarized in Supplementary Table 2. Source data are provided.

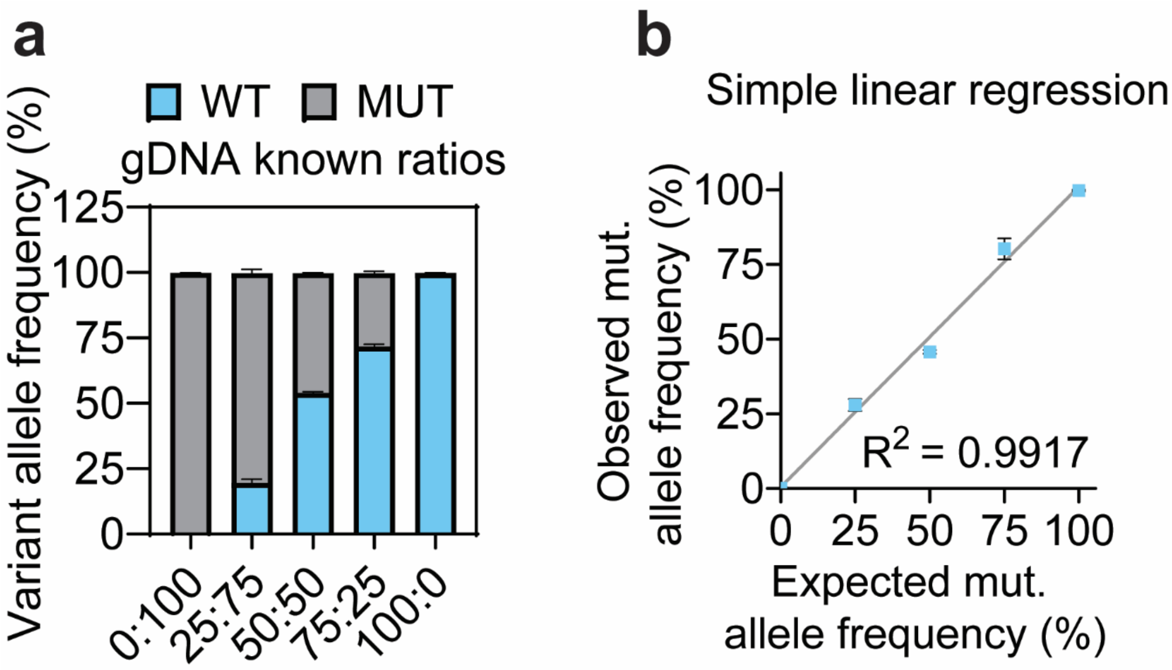


**Supplementary Fig. 4: Validation of k13 variant allele frequency quantification by nanopore sequencing.**

**a,** Known mixtures of wild-type (WT) and mutant (MUT) genomic DNA were prepared at predefined WT:MUT ratios (0:100, 25:75, 50:50, 75:25, and 100:0) and subjected to the targeted k13 amplicon nanopore sequencing pipeline. Genomic DNA was sourced from the parental SenTh15.14 isolate (WT) and a clonal PfKelch13-edited line (MUT). Stacked bars display the mean observed variant allele frequency (%) of WT (blue) and MUT (grey) sequence reads. Error bars indicate standard deviation (s.d.) across independent mixture preparations (n = 3).

**b,** Simple linear regression comparing the expected mutant allele frequency based on the prepared gDNA ratios against the observed mutant allele frequency measured by the nanopore assay. Points represent the mean ± s.d. The highly linear correlation ($R^2 = 0.9917$) demonstrates excellent quantitative accuracy and confirms that the amplicon sequencing pipeline does not introduce significant amplification or sequencing bias toward either allele.

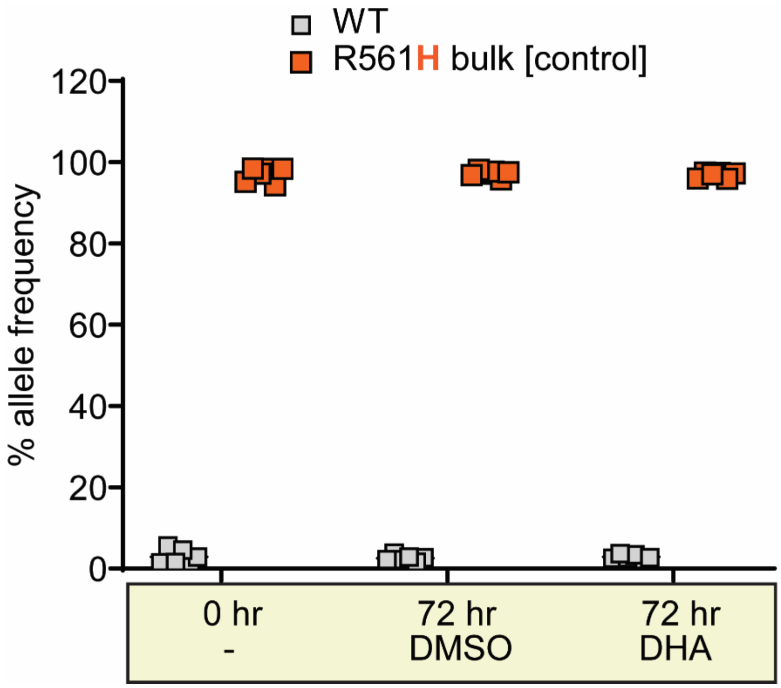


**Supplementary Fig. 5: Allele frequency of a bulk-edited R561H control population remains stable during the nanopore-assisted RSA protocol.**

Variant allele frequency (VAF) of a bulk-edited *Pf*Kelch13 R561H control population was quantified by nanopore sequencing of the *k13* amplicon. Samples were assessed at 0 hr (immediately prior to treatment) and at 72 hr following either a 6 hr 700 nM DHA pulse or a DMSO vehicle control. Grey squares represent the WT allele proportion; orange squares represent the R561H mutant allele proportion. In contrast to the unfit I543T mutation (Fig. 1b), the R561H allele frequency remained highly stable (~97%) throughout the 72-hour assay timeframe. No significant difference was observed between the matched 72 hr DHA and 72 hr DMSO samples (P = 0.585, two-tailed paired t-test), confirming that the nanopore-assisted RSA protocol does not introduce artifactual shifts in allele frequency for stable *k13* variants.

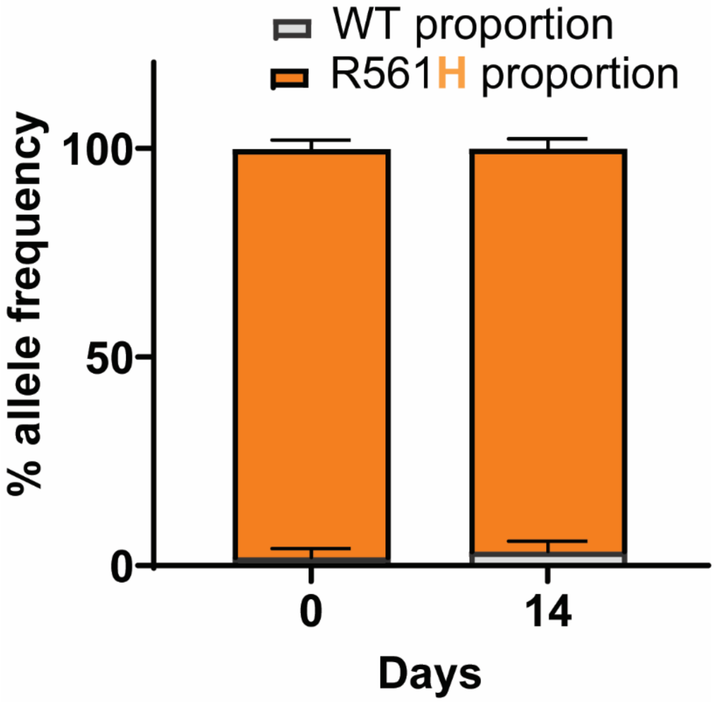


**Supplementary Fig. 6: Allele frequency of a bulk-edited R561H control population remains completely stable during 14 days of drug-free culture.**

Drug-free bulk culture dynamics of a *Pf*Kelch13 R561H control population. Bulk-edited cultures carrying the R561H mutation were maintained under standard *in vitro* conditions without drug selection and sampled at day 0 and day 14. WT (grey) and mutant (R561H, orange) variant allele frequencies (VAFs) were quantified by k13 amplicon nanopore sequencing. Stacked bars show the mean proportion (WT + mutant = 100%) ± s.d. across biologically independent bulk cultures (n = 2). In stark contrast to the severe fitness defect and rapid decline observed in bulk-edited I543T populations over the identical 14-day timeframe (Fig. 1d), the R561H allele frequency remained entirely stable. This confirms that the rapid counter-selection of I543T is a mutation-specific biological fitness penalty in the SenTh15.14 background, rather than a general artifact of maintaining bulk-edited mixed cultures over prolonged periods.

**Supplementary Table 1. Primers and gRNA oligonucleotides used for CRISPR-Cas9 editing of *Pfkelch13.*** Sequences are shown 5′–3′. Donor templates were generated using approximately 500 bp 5′ (upstream) and 3′ (downstream) homology arms flanking the target codon; 5′ and 3′ are defined relative to the *Pfkelch13* 3D7 reference sequence used for donor design. Homology-arm primers include 5′ cloning extensions and *XbaI* and *SalI* restriction sites for insertion into the pUC19 donor backbone. Mutagenesis primers introduce the indicated nonsynonymous SNP (highlighted in red) together with synonymous shield mutations (highlighted in green) to prevent Cas9 re-cutting following HDR-mediated integration. Sequencing-confirmation primers amplify the edited region for Sanger verification. gRNA oligonucleotides are listed as forward/reverse annealing pairs with *BbsI*-compatible overhangs for directional cloning into the Cas9/gRNA expression plasmid.

| Primer ID | Sequence 5'-3' | Use |
|---|---|---|
| K13_5HA1_Fwd | gatcTCTAGAAAGAAGAACATAGGAAACGATTTG | C473S donor: 5' homology arm forward primer |
| K13_3HA1_Rev | atgcGTCGACATTATCAATACCTCCAACAACATATAT | C473S donor: 3' homology arm reverse primer |
| K13_C473S_3HA_Fwd | AGTACTCCGATGTCAACGAAGAAAGCTTATTTTGGAAGTGCTGTAT | C473S: edit-site (mutagenesis) forward primer |
| K13_C473S_5HA_Rev | TCTTCGTTGACATCGGAGTACTCATACGCCAGCATTGTTGACT | C473S: edit-site (mutagenesis) reverse primer |
| K13_5HA2_Fwd | atgcTCTAGACTTAGATAGGGATAGTGAGTTATT | 543/561/579 donors: 5' homology arm forward primer |
| K13_3HA2_Rev | atgcGTCGACTTATATATTTGCTATTAAAACGGAGTGA | 543/561/579 donors: 3' homology arm reverse primer |
| K13_I543T_3HA_Fwd | AGGATCTATTGTACAGGGGGATATGATGGCTCTTC | I543T: edit-site (mutagenesis) forward primer |
| K13_I543T_5HA_Rev | TCCCCCTGTACAATAGATCCTACCATTTGACGTAACACCAC | I543T: edit-site (mutagenesis) reverse primer |
| K13_R561H_3HA_Fwd | TCATCATATGAAGGCCTGGGTAGAGGTGGCACCTT | R561H: edit-site (mutagenesis) forward primer |
| K13_R561H_5HA_Rev | CCCAGGCCTTCATATGATGATCATATGCTTCTACATTC | R561H: edit-site (mutagenesis) reverse primer |
| K13_M579I_3HA_Fwd | TCCAAGATCTTCTGCAATTTGTGTTGCTTTTGATAATAAAATTTATGT | M579I: edit-site (mutagenesis) forward primer |
| K13_M579I_5HA_Rev | ACACAAATTGCAGAAGATCTTGGAGTATTCAAAGGTGCCACCTCTA | M579I: edit-site (mutagenesis) reverse primer |

| | | |
|---|---|---|
| K13_5HA3_Fwd | CTTAGATAGGGATAGTGAGTTATTTA G | Sequencing confirmation forward primer |
| K13_3HA3_Rev | GAACATAACATATTAGATTCCGTTGAA C | Sequencing confirmation reverse primer |
| **Selected guides for gRNA insertion into Cas9 plasmids for *Pfkelch13* gene editing** | | |
| K13_gRNA1_Fwd | TATTGTAAGCTTTTTTGGTAGACAT | C473S gRNA oligo (forward) |
| K13_gRNA1_Rev | AAACATGTCTACCAAAAAAGCTTAC | C473S gRNA oligo (reverse) |
| K13_gRNA2_Fwd | TATTGGGTAGAATTTATTGTATTGG | I543T gRNA oligo (forward) |
| K13_gRNA2_Rev | AAACCCAATACAATAAATTCTACCC | I543T gRNA oligo (reverse) |
| K13_gRNA3_Fwd | TATTGTGATCATCGTATGAAAGCAT | R561H gRNA oligo (forward) |
| K13_gRNA3_Rev | AAACATGCTTTCATACGATGATCAC | R561H gRNA oligo (reverse) |
| K13_gRNA4_Fwd | TATTGACACATAGCTGATGATCTAG | M579I gRNA oligo (forward) |
| K13_gRNA4_Rev | AAACCTAGATCATCAGCTATGTGTC | M579I gRNA oligo (reverse) |

**Supplementary Table 2: *In vitro* drug susceptibility of the SenTH15.14 parent line and isogenic *Pfkelch13*-edited derivatives.**

Half-maximal effective concentration ($EC_{50}$) values were determined for dihydroartemisinin (DHA), chloroquine (CQ), monodesethylamodiaquine (mdADQ), lumefantrine (LUM), and piperaquine (PPQ) in the SenTH15.14 WT parent line and CRISPR-engineered *Pfkelch13* mutant lines (R561H, I543T, M579I, and C473S) generated in the same genetic background. Values are presented as mean ± SD from three independent biological replicates. $EC_{50}$ values for the CRISPR-edited lines did not differ significantly from those of the SenTH15.14 WT parent for the tested compounds. Dose–response ($EC_{50}$) curves for each drug and parasite line are shown in **Supplementary Fig. 3**.

| | DHA | CQ | mdADQ | LUM | PPQ |
|---|---|---|---|---|---|
| *Parent (WT)* | 1.63 ± 0.2 | 76.9 ± 25.8 | 18.0 ± 5.3 | 1.08 ± 0.3 | 4.66 ± 1.0 |

| | | | | | |
|---|---|---|---|---|---|
| *R561H* | 1.01 ± 0.6 | 82.7 ± 24.5 | 15.8 ± 2.9 | 0.80 ± 0.2 | 5.43 ± 1.1 |
| *I543T* | 1.07 ± 0.6 | 80.7 ± 17.5 | 18.3 ± 4.4 | 1.12 ± 0.1 | 5.45 ± 0.8 |
| *M579I* | 1.12 ± 0.2 | 94.5 ± 7.5 | 23.5 ± 0.2 | 0.89 ± 0.1 | 6.26 ± 0.4 |
| *C473S* | 1.06 ± 0.6 | 91.7 ± 34.4 | 19.5 ± 6.1 | 0.74 ± 0.1 | 4.90 ± 0.7 |
